# Systematic inference of biological relatedness across the global ancient DNA record

**DOI:** 10.64898/2026.11.19.767913

**Authors:** Edward C. Huang, Harald Ringbauer

## Abstract

The rapidly expanding scale of ancient DNA data creates an increasing need for high-performance software methods. One area for such improvement is the inference of pairwise biological relatedness, a prominent aspect of many ancient DNA studies. We present `fastpmr`, a fast and memory-efficient method for biological relatedness inference based on the computation of pairwise mismatch rates. We demonstrate that `fastpmr` can achieve over 1000× speed-up over `READv2`, a widely used relatedness-inference method. We leverage this performance to investigate relatedness at the scale of the global ancient DNA record. Analyzing biological relatedness across all pairs in the Allen Ancient DNA Resource (*n* = 19,118 ancient and 23,089 total samples) recapitulates human evolutionary history and identifies previously unknown genetic duplicates and likely sample mix-ups, demonstrating `fastpmr`’s utility for ancient DNA analysis and quality control at scale.

## 1 Introduction

In recent years, ancient DNA (aDNA) datasets have expanded rapidly [1]; at the time of writing, the Allen Ancient DNA Resource (AADR) includes 19,118 ancient samples [2]. This increase in scale has enabled novel analyses of the human past, including of population history [3–5, etc.], paleoepidemiology [6, 7], and natural selection [8–14, etc.].

As aDNA datasets continue to grow, new software methods are needed to handle these data efficiently at scale. One important component of aDNA analysis is the inference of biological relatedness. Beyond its direct importance in inferring social and kinship practices from aDNA [15, 16], biological relatedness information is essential to both ancient and modern DNA analyses, as closely related individuals can distort genetic inference and downstream analyses. Additionally, identifying genetically identical samples is a crucial quality-control step in aDNA analysis, as it reveals individuals who were unknowingly sampled repeatedly (as is common with commingled human remains) or sample mix-ups in the data pipeline, including archaeological curation, wet lab analysis, and bioinformatic processing. Here, we define genetically identical samples as samples originating from identical genomes, whether from one individual or from identical twins. Finally, biological relatedness information enables inference of biological connectivity across time and space, furthering our understanding of evolutionary and population history.

A commonly used measure of biological relatedness in aDNA is the pairwise mismatch rate (PMR), which refers to the average allelic mismatch rate between two genetic sequences, computed over all autosomal sites (see Methods: PMR computation) [17]. The PMR is closely related to other measures of genetic diversity, such as nucleotide diversity (*π*) (see Results: Inference of human evolutionary history). Many existing software methods compute PMRs for aDNA datasets and then use these PMRs to classify degree-level kinship relationships between individuals, up to the second or third degree [17–21]. However, these methods are used primarily for relatively small aDNA datasets and are not optimized to infer biological relatedness across thousands or tens of thousands of individuals. Another commonly used measure of biological relatedness is identity-by-descent (IBD) segment sharing. IBD-sharing information, which reflects the exact genomic segments co-inherited from recent shared ancestors, is a higher-resolution metric of biological relatedness and can detect more distant relatives up to around the sixth degree [22]. However, compared to PMR-based methods, IBD inference requires substantially higher-coverage samples, which limits its applicability [18, 22].

To address the lack of methods for inferring biological relatedness in large aDNA datasets, we present `fastpmr`, a `Rust` package that efficiently computes PMRs between ancient individuals (see Methods: Software implementation). `fastpmr` enables fast and memory-efficient inference of biological relatedness in large aDNA datasets, supporting ever-larger studies of ancient history, connectivity, and evolution. In our benchmarking experiments, we find that `fastpmr` runs over 1,000 times faster than `READv2` [18], a commonly used method for ancient relatedness inference, while preserving `READv2`’s PMR and degrees-of-relatedness results. This performance enables us to use `fastpmr` to compute PMRs across the entire AADR [1, 2], which contains 19,118 ancient samples and is, to our knowledge, the largest publicly available aDNA dataset. In doing so, we show that ancient biological relatedness information reflects human evolutionary history, including the Out of Africa migration and the peopling of the Americas from Asia. Additionally, we find that background relatedness has decreased significantly over the past 10,000 years in Europe and Asia, but not significantly in the Americas. Finally, we use `fastpmr` to identify potential errors and cryptic duplicates in the global aDNA record, demonstrating the utility of large-scale PMR screens enabled by `fastpmr`.

## 2 Results

### 2.1 Performance benchmarks

We conducted several benchmarks to evaluate `fastpmr`’s computational resource requirements. We first compare `fastpmr`’s performance to that of `READv2` [18], a commonly used method for inferring ancient kinship. We run both methods on an ancient DNA dataset of individuals from the Southern Cone of South America, comprising 238 samples and *∼*1.9 million autosomal variants [23, 24]. We refer to this dataset as the “Maravall-López dataset.” We find that `fastpmr` represents an extremely performant method across various sample sizes (Figure 1, see Methods: Software implementation). `fastpmr` is designed to exactly reproduce `READv2`’s PMR and degrees-of-relatedness results. For the `fastpmr` and `READv2` results on the Maravall-López dataset, all 28,203 PMRs agree within a tolerance of 10^*−*6^, and all degrees-of-relatedness classifications are identical. However, we note that `READv2` infers some additional information beyond PMRs, including first-degree relation types (parent–child vs. sibling) and block-jackknife standard errors.

**Figure 1:**
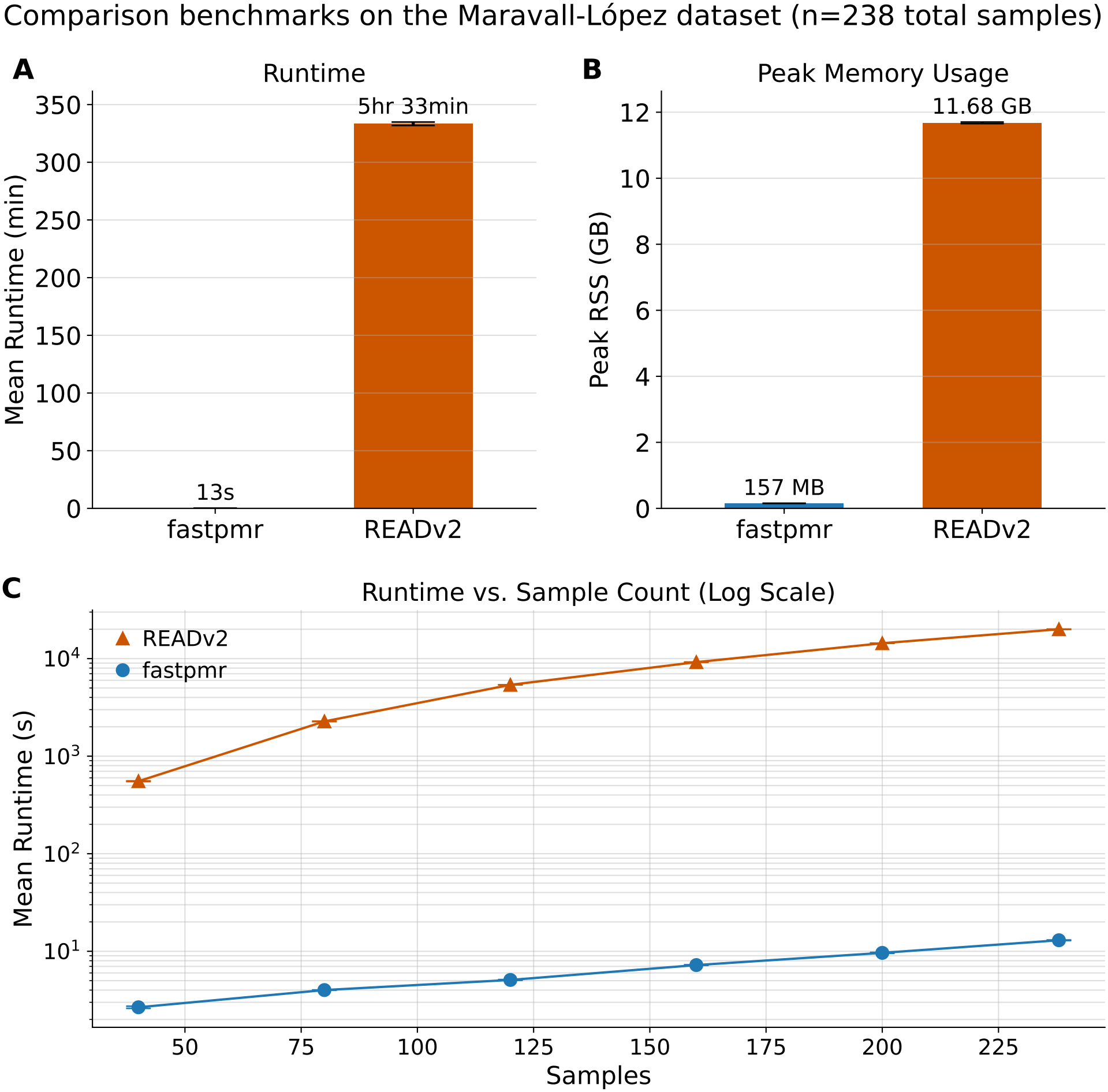
Performance comparison of `fastpmr` and `READv2`, showing (**A**) mean runtime, (**B**) mean peak memory usage, and (**C**) mean runtime vs. sample count on the Maravall-López dataset. Error bars denote the standard deviation of three runs.

We also evaluate the effect of several variables on `fastpmr` runtime and memory usage (Figure 2). We run these benchmarks using an ancient DNA dataset of individuals from across Eurasia, comprising 1,383 samples and *∼*1.15 million autosomal variants [25, 26]. We refer to this dataset as the “Lazaridis dataset.” For benchmarks on the Lazaridis dataset, sample, variant, and thread counts are set using `fastpmr`’s relevant command-line flags (Methods: Software implementation). Each benchmark on the Lazaridis and Maravall-López datasets is run on an exclusively provisioned compute node of the “GRACE” HPC cluster at the Max Planck Institute for Evolutionary Anthropology; each of these nodes is equipped with two AMD EPYC 9745 CPUs.

**Figure 2:**
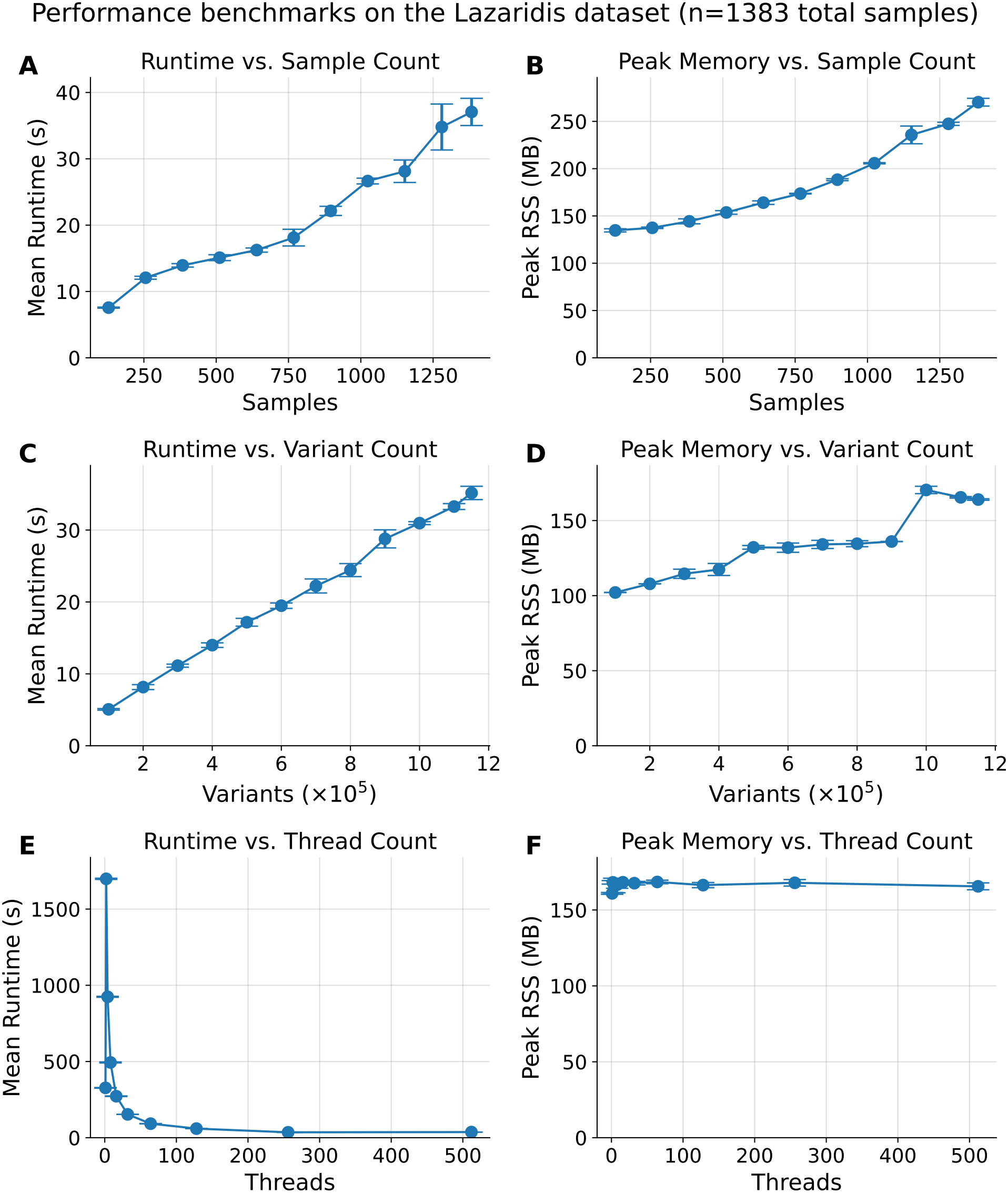
Performance of `fastpmr` with varying (**A, B**) sample counts, (**C, D**) variant counts, and (**E, F**) thread counts on the Lazaridis dataset [25, 26]. Error bars denote the standard deviation of three runs.

First, we examine the effect of sample count on `fastpmr` performance. Subsetting the Lazaridis dataset, we create sample sets of varying size. We then run `fastpmr` on the Lazaridis dataset while limiting PMR computations to these sample subsets. Since the number of pairwise comparisons *n*(*n −* 1)*/*2 scales asymptotically quadratically with the sample count *n*, we expect both runtime and peak memory usage to scale quadratically with large *n*. However, we observe a more linear relationship between sample count and performance for sample counts up to 1250, suggesting that computational overhead plays a significant role at these sample counts (Figure 2A,B).

To also examine the effect of variant count on `fastpmr` performance, we run `fastpmr` on the Lazaridis dataset while varying the number of included genetic variants. We find that runtime appears to increase linearly with variant count as expected, while peak memory usage increases only marginally with variant count (Figure 2C,D).

Finally, we examine the effect of thread count on `fastpmr` performance. In this benchmark, we run `fastpmr` on the Lazaridis dataset while varying its thread-count parameter. We find that while peak memory usage remains relatively constant for all thread counts, runtime first increases sharply when switching from single-threaded to multi-threaded usage before then decreasing at higher thread counts (Figure 2E,F). These results appear to reflect the computational overhead incurred in multi-threaded usage; the runtime benefit of multi-threading does not outweigh this overhead at lower (non-single) thread counts. Specifically, multi-threading in `fastpmr` introduces overhead from dispatching computation across threads and atomically incrementing shared result arrays. In this benchmark, we begin to see runtime benefits from multi-threading at around 16 threads.

### 2.2 Inference of human evolutionary history

We run `fastpmr` on the 1240k dataset of AADR v66.p1, which contains 19,118 ancient samples and 23,089 total samples. Unless otherwise specified, we refer to this specific dataset as the “AADR.” Here, we compute the `fastpmr`-computed PMRs of all *∼*267 million sample pairs in the AADR and analyze various subsets of these PMR data. Note that we refer to AADR entries as “samples” instead of “individuals” as multiple entries may originate from the same individual. Computing all PMRs in the AADR took 2.81 hours with a peak memory usage of 31.8 GB on an exclusively provisioned compute node of the “GRACE” HPC cluster at the Max Planck Institute for Evolutionary Anthropology (see Results: Performance benchmarks).

The median PMR across all pairs in a group is commonly used as a measure of background relatedness [18]; higher median PMRs indicate lower background relatedness, and vice versa. PMR is also closely related to other measures of genetic diversity. For example, the PMR between two haploid genetic sequences is exactly their pairwise nucleotide diversity (*π*). Note that these metrics are subject to ascertainment over a given SNP set, which for aDNA data is often the 1240k SNP set. Additionally, although population-wide nucleotide diversity is defined as a mean value, median PMR values are widely used in aDNA analyses because the median is more robust than the mean to bias from close kinship relationships. As such, we use the median PMR of an AADR locality (for a given time transect) as a proxy for genetic diversity. Throughout this work, we retrieve AADR localities—which generally correspond to archaeological sites—directly from the AADR metadata.

Here, we report and analyze median within-locality PMRs in the AADR, providing insights into the dynamics of genetic diversity across space and time. Notably, we find that this ancient biological relatedness data recapitulate consensus paths of human migration, including the Out of Africa event and the peopling of the Americas from Asia (Figure 3). Specifically, background relatedness remains consistently low in Europe and increases along a gradient corresponding to anatomically modern humans’ migration paths east across Eurasia and into the Americas, where localities show consistently higher background relatedness.

**Figure 3:**
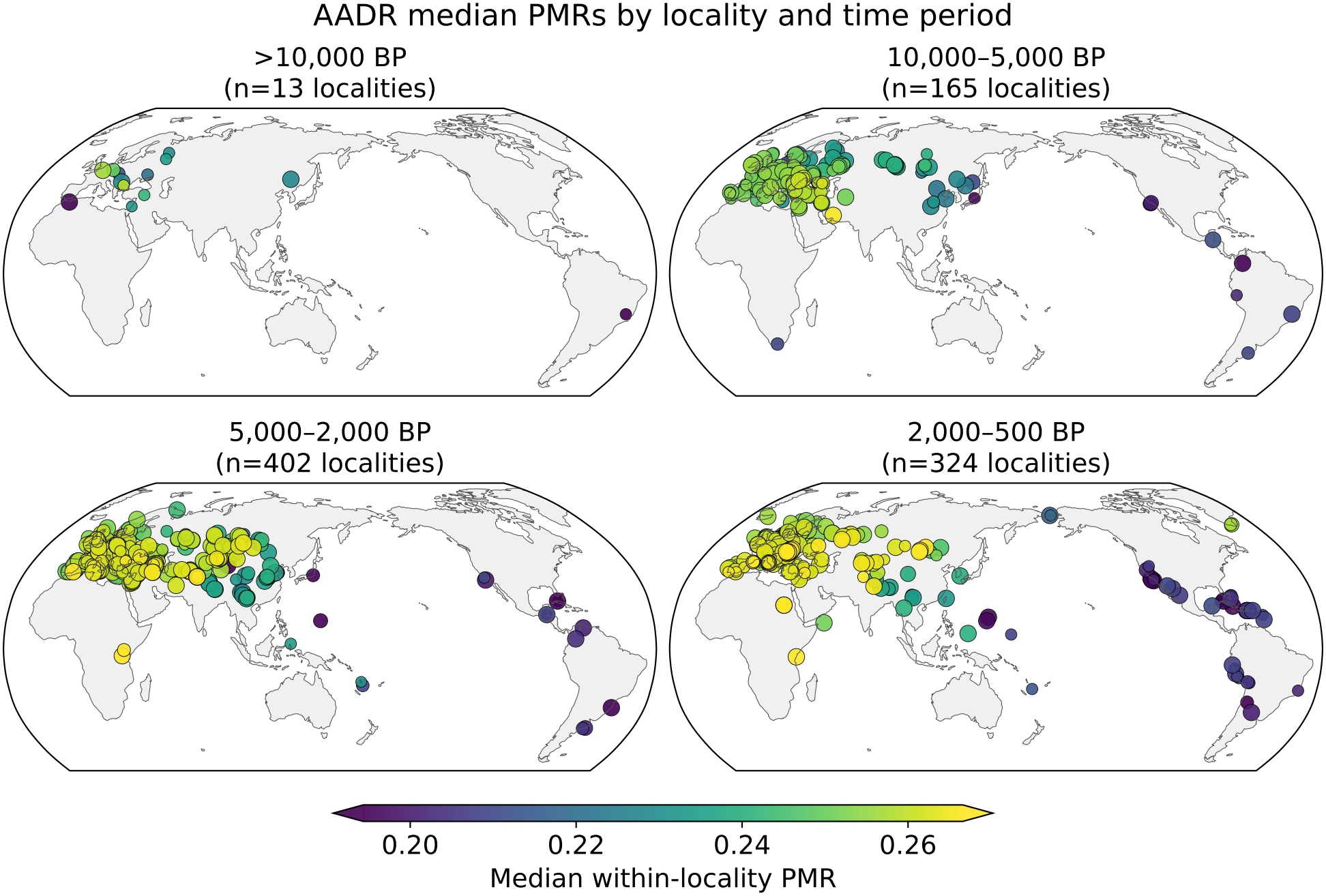
Median pairwise mismatch rates (PMRs) of localities in the Allen Ancient DNA Resource (AADR), plotted over space and time. Each point represents the median PMR of a locality with at least five qualifying sample pairs in the AADR for a given time period. Only sample pairs with differing AADR individual IDs and at least 30,000 overlapping sites are considered. Point size scales with the number of qualifying sample pairs, up to 20 pairs.

To further investigate the relationship between geography and genetic diversity, we plot within-locality PMR against migratory “Out of Africa” distance (Figure 4). To calculate migratory distance, we approximate the approach of Ramachandran et al. [27] by computing distance from Addis Ababa via predetermined waypoints, assuming that most human migrations did not traverse large bodies of water. We observe a strong negative relationship between median within-locality PMR and “Out of Africa” migratory distance. Additionally, outlier localities on this scatterplot are readily explainable, including a group of medieval individuals with Ashkenazi Jewish-like ancestry that underwent a strong population founder effect, and a pair of Norse Greenland settlements that circumvent Greenland’s waypoint-prescribed migratory path through the Americas. We also note that this scatterplot appears to show many more low-PMR outliers than high-PMR outliers; this pattern likely reflects the relative rarity of events that increase genetic diversity (e.g., admixture) compared with those that reduce it (e.g., genetic drift and population founder effects).

**Figure 4:**
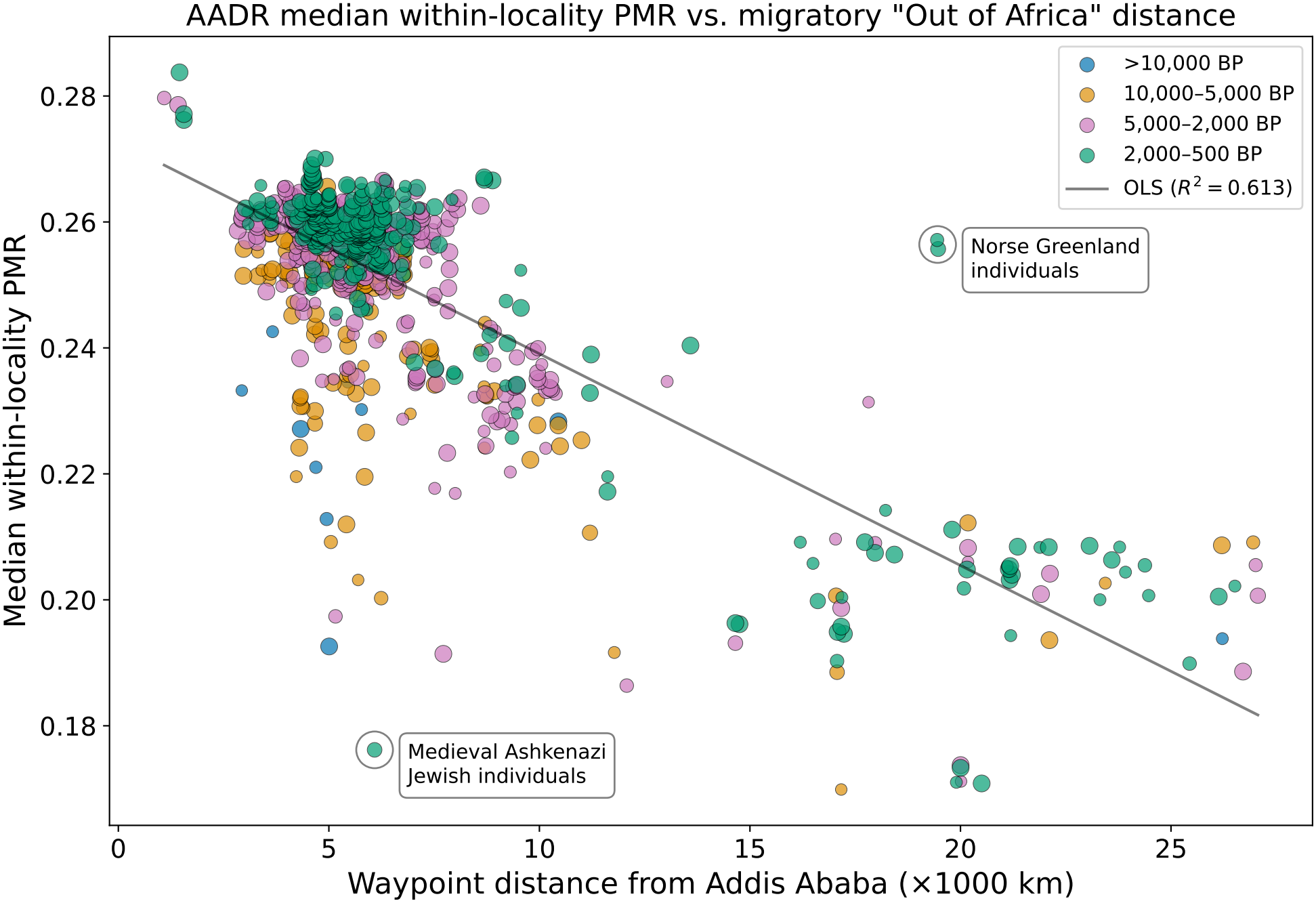
Median pairwise mismatch rates (PMRs) of localities in the Allen Ancient DNA Resource (AADR), plotted against “migratory” waypoint distance from Addis Ababa, Ethiopia. Each point represents the median PMR of a locality with at least five qualifying sample pairs in the AADR from a given time period. Only sample pairs with differing AADR individual IDs and at least 30,000 overlapping sites are considered. Point size scales with the number of qualifying sample pairs, up to a limit of 20 pairs. The regression line is computed with ordinary least-squares, and selected outliers are annotated.

We also plot the change in genetic diversity over time for different global regions (Figure 5). We find that genetic diversity has significantly increased over time in Europe and Asia, but not in the Americas. Additionally, genetic diversity in Europe and Asia remains consistently higher than in the Americas from 10,000–5,000 BP. The spread of median within-locality PMRs in Europe also appears generally smaller than in Asia and the Americas.

**Figure 5:**
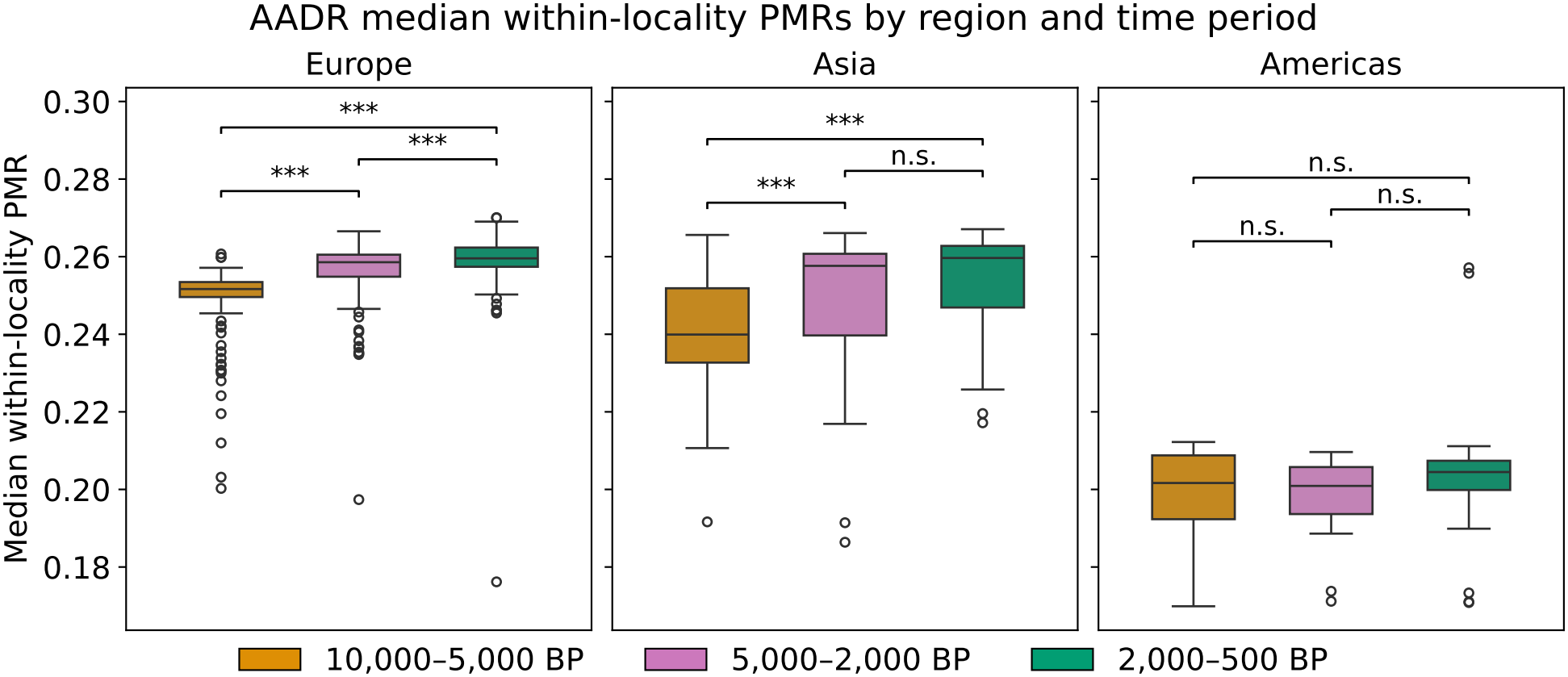
Median pairwise mismatch rates (PMRs) of localities in the Allen Ancient DNA Resource (AADR), stratified by region and time period. Localities with at least five qualifying sample pairs in the AADR for a given time period are included. Only sample pairs with differing AADR individual IDs and at least 30,000 overlapping sites are considered. Each box spans the 25th to 75th percentile of PMRs, and whiskers extend past box edges with maximum length 1.5 × IQR (interquartile range). Brackets show significance levels of pairwise Mann-Whitney U tests between time periods, with Holm correction applied across all nine comparisons (***: *p <* 0.001, **: *p <* 0.01, *: *p <* 0.05, n.s.: *p ≥* 0.05).

Finally, we conduct an exploratory analysis leveraging between-locality PMRs in the AADR, training a Gaussian process model to predict PMRs given pairs of space-time coordinates (see Supplementary Note 1: Gaussian process analysis).

### 2.3 Quality control and discovery at scale in the global aDNA record

The AADR is a regularly updated and widely used repository of aDNA data. To demonstrate how `fastpmr` enables the identification of potential data processing issues and unknown kinship connections in the global aDNA record, we use `fastpmr`-computed PMRs to run various large-scale “scans” over the AADR (Supplementary Tables S1-3).

First, we identify pairs of samples labeled in the AADR as originating from the same individual but sharing high PMR values inconsistent with being genetic duplicates (PMR *>* 0.17)(Supplementary Figure S3A). We refer to these pairs as “potential spurious duplicates.” We find 40 such sample pairs, which are reported in Supplementary Table S1: AADR v66.p1 Potential Spurious Duplicates. Notably, only three (purported) individuals appear in 30 of these sample pairs, suggesting data-processing issues for these three individuals.

Next, we identify pairs of samples labeled in the AADR as originating from different individuals but sharing low PMR values consistent with being genetic duplicates (PMR *<* 0.14)(Supplementary Figure S3B). We refer to these sample pairs as “potential cryptic duplicates.” We identify 129 such sample pairs, reported in Supplementary Table S2: AADR v66.p1 Potential Cryptic Duplicates (Annotated). We also annotate apparent characteristics of these sample pairs (Table 1); over half of the sample pairs share at least one similar AADR metadata field, strongly indicating that they are true duplicates originating from the same individual. We further highlight three particularly noteworthy cases of potential cryptic duplicates below.

**Table 1:** Characteristics of 129 potential cryptic duplicate pairs in the AADR. Categories are not necessarily mutually exclusive, so counts do not sum to the total. Pairs with similar AADR IDs (such that they clearly represent duplicates) are not evaluated for other categories besides same/different source publication. All annotations are made to the best of the authors’ knowledge and may not be comprehensive. “Pub.”: publication.

| Category | *N* |
|---|---|
| Bioinformatically identical sample (PMR = 0) | 20 |
| Similar AADR ID | 21 |
| Similar AADR skeletal code | 58 |
| Flagged in source pub. as genetically identical | 10 |
| Flagged in source pub. as genetically non-identical | 6 |
| Flagged in AADR metadata | 3 |
| Different source pub. | 79 |

First, our “cryptic duplicate” scan flags a pair of samples from a ritual site at the ancient settlement of Chichén Itzá: YCH009 and YCH049, which were previously described in Barquera et al., 2024 [28]. Notably, the original study identifies two other pairs of identical twins interred at the same site — each with PMRs of *∼*0.11 — interpreted as evidence of a preference for close biological kinship in Maya ritual practice. These two pairs do not include YCH009–YCH049; the original study infers this specific pair to be unrelated, with a reported PMR of *∼*0.20. However, our scan reveals a much lower PMR of *∼*0.12 for YCH009–YCH049. To confirm this deviating result, we then recompute PMRs for the Chichén Itzá samples using genotypes called directly from the original study’s published sequence data (Supplementary Table S4: Chichén Itzá Recomputed PMRs). In this analysis, we infer a YCH009–YCH049 PMR of *∼*0.12 and a within-site median PMR of *∼*0.21, which is again concordant with YCH009 and YCH049 being identical twins. In both analyses, YCH009 and YCH049 share over 250,000 overlapping sites, which rules out classification uncertainty due to low genome coverage. In summary, this scan potentially identifies a third pair of identical twins from a ritual site at Chichén Itzá, further highlighting the apparent emphasis on close biological kinship in Maya ritual practice. An alternative explanation is that a data mix-up occurred before the original study’s PMR computation and before the sequence data were deposited. We focus here on YCH009–YCH049, but we note that several other non-twin kinship relationships reported in the original study are not reproduced in the AADR or published sequence data. These relationships include YCH034–YCH041, YCH049–YCH057, and YCH047–YCH057, which were originally reported as close relative pairs but show no evidence of close kinship in our analyses. We contacted the authors of the original Chichén Itzá study regarding these findings, prompting investigation into the apparent data discrepancy. (We note that YCH049 appears to be mislabeled in AADR v66.p1 as “YCH017 alt1”, and that we recover the apparent true label of YCH049 from the entry’s skeletal code. The sample pair YCH009–YCH049, labeled as such, is also identified as a potential cryptic duplicate in the AADR v62.0.p1 scan mentioned below, and the PMR for this sample pair remains similar when computed from the original study’s published sequence data, as mentioned above.)

Second, the “cryptic duplicate” scan flags three samples with Ashkenazi Jewish-like ancestry: SB605, SB606, and SB671, representing three of the six individuals originally reported in Brace et al., 2022 [29]. These samples, originating from the same locality in medieval England, were reported to belong to a trio of full-sibling sisters. However, in our PMR data, the sample pairs SB605–SB606 and SB605–SB671 each share a PMR of around 0.09, measured on over 100,000 overlapping sites; the remaining pair SB606–SB671 shares a PMR of 0.067 over 25,509 overlapping sites, which did not cross the original scan’s site-overlap threshold. Given the locality’s median PMR value of 0.176, the PMR values for SB605, SB606, and SB671, which are about half of the locality’s median PMR (rather than the 75% of median expected for first-degree relatives), suggest that these three samples are genetically identical. All three samples originate from genetic females [29], consistent with their being genetically identical. Furthermore, these samples were osteologically inferred to originate from a 5-10 year old, a 10-15 year old, and a “young adult”—three similar ages—and all three samples derive from three different anatomical sources (petrous bone, molar, and femur) [29]. Finally, this locality’s archaeological context is a commingled mass burial, which increases the likelihood that these samples derive from the same individual. We confirm the three samples to be genetically identical by using an independent line of evidence. We infer runs of homozygosity (ROH) for each sample using the software `hapROH` [30] with a custom reference panel optimized for low-coverage whole-genome sequencing data. This analysis shows that each sample’s ROH distribution is almost identical, a pattern that would be extremely unlikely if the three samples belonged to full siblings (see Supplementary Note 2: ROH analysis of samples from medieval England).

Additionally, in a similar `fastpmr` scan of a previous AADR release (v62.0), we identified four potential cryptic duplicates where the sample pair spanned two different studies published in Nature on the same day in 2019 [31, 32]. Moreover, a reanalysis of samples from one of these original studies [33] yielded three more potential cryptic duplicates with the other original study. These sample pairs, all from Siberia, highlight the possibility that aDNA data can be unknowingly duplicated across studies. We notified AADR maintainers of these potential genetic duplicates, most of which were updated in AADR v66.p1. We originally detected these potential errors using earlier versions of `fastpmr` and our scan pipeline. We report updated AADR v62.0.p1 [2] scans in Supplementary Tables S5-7. We rerun these scans with updated versions of `fastpmr`, our scan pipeline, and AADR v62.0 (v62.0.p1, which removes certain modern samples), but they recover the same potential cryptic duplicates described here, plus one additional pair from the two 2019 studies.

In our final empirical application, we aim to identify biological relatedness between samples from different localities in the AADR. Since inference of close relatives is more sensitive to variation in region-specific baseline PMR values compared to inference of identical individuals, we limit this scan to sample pairs from Eurasia. Eurasian samples comprise 19,122 of the 23,089 samples in the AADR.

We set the PMR threshold for this scan at 0.14 *≤* PMR *<* 0.18, which generally identifies close kinship (1st-degree) in Eurasian sample pairs. We find 40 such sample pairs, which are reported in Supplementary Table S3: AADR v66.p1 Potential Cross-Site Relatives. However, certain samples are overrepresented in this list beyond what is biologically plausible, suggesting a data-generation or data-processing error that systematically deflates these samples’ PMRs. For example, severe reference bias in read alignment can cause a sample to preferentially drop alternate-allele reads, shifting its genotypes toward the reference allele. Since the reference allele is usually also the major allele, such a sample would mismatch all other samples at a reduced rate, producing spuriously low PMRs. Detecting such samples is an important quality control step for aDNA datasets.

We identify one particularly notable case of long-distance close kinship, between two individuals sampled approximately 1,800 kilometers apart in Kazakhstan and first reported in Gnecchi-Ruscone et al., 2021 [34]. However, these two individuals, BSB004 and KLK002, are associated with very different dates based on their respective archaeological contexts: BSB004 is dated to 500–200 BCE, while KLK002 is dated to 800–1100 CE. Originally, neither sample was directly dated. Given BSB004’s close kinship with KLK002—which is virtually impossible over a 1000-year gap—and the context date of the Bisoba burial ground (500–200 BCE), where BSB004 was sampled, we originally hypothesized two plausible explanations. The first was that BSB004 represents a sample mix-up, and the second was that BSB004 represents a genuine later burial at the Bisoba burial ground, which would be a striking case of medieval long-distance mobility. Based on this finding, follow-up analysis of these individuals was initiated, including direct radiocarbon dating of each individual’s archaeological sample and Twist capture of the original BSB004 library. Carbon-14 dating results, which returned before submission of this work, confirm the previously inferred *>*1000-year gap between the archaeological samples (see Supplementary Note 3: Radiocarbon dating of samples from Kazakhstan), precluding the possibility that BSB004 represents a later burial at Bisoba and instead suggesting a sample mix-up or contamination. Sequencing of the Twist-captured BSB004 library revealed that most reads carried the index pair of a KLK002 library, indicating a laboratory mix-up. Altogether, our results show that screening for biological relatives is an important aDNA quality-control step in itself.

## 3 Discussion

As the scale of aDNA data continues to increase, aDNA software methods must adapt to keep pace. The inference of biological relatedness is an especially important methodological domain, as its runtime generally scales quadratically with the number of samples. We find that `fastpmr` represents a significant performance advance in this domain, enabling analyses of ancient relatedness and connectivity at unprecedented scale. Analysis of the AADR with `fastpmr` reveals global patterns of human evolution, and we believe `fastpmr` generally facilitates larger studies of aDNA data that will advance our understanding of the past. We note that our analysis of AADR PMRs reflects period-specific insights into (pre)historic individuals and migration events, which are not necessarily relevant to present-day populations. Additionally, the relative scarcity of aDNA data from certain world regions, such as Africa and the Americas, further limits the scope of our analysis.

We also find that `fastpmr`’s performance enables improved quality control for large aDNA datasets, identifying both potential errors and unknown duplicates across studies. Such quality control at scale will be crucial as the global aDNA record continues to expand and mature. Another potential application of `fastpmr` is in the inference of “baseline” PMRs. While the median PMR of a locality is often used as a proxy for the expected PMR of two unrelated individuals from that locality, this approach may be inappropriate for localities with diverse ancestries or for datasets encompassing multiple localities. In this case, large-scale PMR data could be used to predict an “expected” PMR for two unrelated samples given their space-time and ancestry data (see Supplementary Note 1: Gaussian process analysis).

Compared with other methods for inferring biological relatedness in aDNA, `fastpmr` offers a significantly more performant alternative and is especially suitable for large-scale analyses. However, other methods present benefits in different aspects of analysis. For example, `BREADR` [21] uses a Bayesian framework for relatedness inference, enabling the output of posterior probabilities. In addition, methods such as `KIN` [35] and `READv2` [18] can classify first-degree kinship relationships as parent–child or full sibling relationships. In summary, existing methods may offer advantages for more detailed analysis of biological relatedness, while `fastpmr` offers advantages for efficient, large-scale analyses of biological relatedness.

In the future, we hope to improve `fastpmr` by incorporating additional functionality present in related methods. For example, in principle, it is possible to classify first-degree relationship types (parent–child vs. full-sibling) and compute bootstrapped estimates of PMR standard errors. However, both features require storing spatial PMR information, so implementation would require reworking `fastpmr`’s internal data structures. As such, care must be taken to add functionality while preserving `fastpmr`’s performance. Ultimately, we believe that this work is only a first step in inferring ancient relatedness at scale and, more broadly, conducting big-data analyses of aDNA.

## 4 Methods

### 4.1 PMR computation

For a pair of individuals *i* and *j*, the PMR (pairwise mismatch rate) is defined as the mean allelic mismatch rate across all autosomal sites (chromosomes 1–22 for humans) at which both individuals have non-missing genotypes. Non-missing genotypes can be encoded as reference allele dosages *g ∈ {*0, 1, 2*}*, corresponding to homozygous alternate, heterozygous, and homozygous reference genotypes, respectively. At each site *s*, the per-site mismatch rate between two individuals *i* and *j* is then

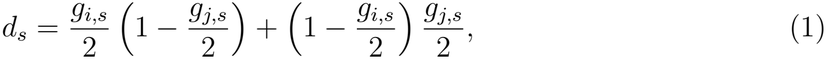

where *g*_*i,s*_ and *g*_*j,s*_ are the genotypes of individuals *i* and *j* at site *s*. This equation represents the probability of drawing mismatched alleles when sampling one allele from each individual. We note that this formulation naturally recovers the expected mismatch between two heterozygous sites (50% mismatch probability).

Given each per-site mismatch rate *d*_*s*_, the PMR between individuals *i* and *j* is then the average over sites

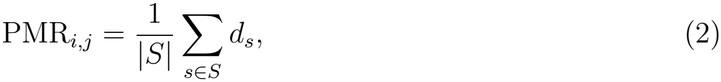

where *S* is the set of autosomal sites at which both individuals have non-missing genotypes [36]. Importantly, this framework also yields correct PMRs for “pseudo-haploid” [1] sequences widely used for aDNA data, where one read is randomly sampled for each variant [37]. In this format, alternate genotypes are represented with a reference allele dosage of 0 and reference genotypes with a reference allele dosage of 2.

### 4.2 Software implementation

`fastpmr` is implemented in `Rust`. It operates in two passes over the input genotype data. In the first pass, `fastpmr` counts the number of non-missing genotypes (covered SNPs) per individual and reports these covered SNP counts. Individuals with fewer than the user-specified minimum number of covered SNPs (default: 30,000) are excluded from subsequent computations. In the second pass, for each site in the input genotype data, `fastpmr` computes *doubled* per-site PMRs for each pair of individuals with non-missing genotypes at that site. Since per-site PMRs can take a value of 0.5, it computes doubled per-site mismatch rates to avoid using floating-point numbers. Doubled per-site PMRs are computed using a lookup table indexed by genotype, and results are accumulated into a flat array storing the sums of these doubled per-site PMRs for each pair of individuals. A corresponding array tracks the doubled count of shared non-missing sites for each pair of individuals. Although the raw count of shared non-missing sites (i.e., the site overlap count) must always be an integer, we still double the count for consistency, ensuring that final PMRs can be computed by dividing the values stored in these two arrays. Both arrays store values as 64-bit unsigned integers, protecting against integer overflow risk.

`fastpmr` supports parallelization of both passes over the input genotype data. When parallelization is enabled, `fastpmr` uses `rayon`’s parallel iterators, which are backed by a work-stealing thread pool [38], to process sites in parallel. Users can specify the number of threads to use; a value of one indicates single-threaded usage. Otherwise, the number of threads used defaults to one for datasets with fewer than 500 individuals and the number of logical cores for datasets with 500 or more individuals.

Input genotype data is read through a unified, iterator-based interface. `fastpmr` supports this interface for `EIGENSTRAT` (unpacked, packed, and transposed packed) and `PLINK` (.bed/.bim/.fam) data formats. `EIGENSTRAT` genotype files are auto-detected as unpacked (plain text), packed (binary, variant-major), or transposed packed (binary, sample-major) based on the .geno file header. `fastpmr` memory-maps transposed packed `EIGENSTRAT` genotype files for efficient access to sites. Otherwise, it streams genotype files from disk.

`fastpmr` presents flexible user options, including the ability to filter input data by individual ID, chromosome, variant index, and covered SNP count, which is computed by `fastpmr` itself. It also optionally classifies degrees of relatedness using computed PMR information. Its degree classification procedure replicates that of `READv2` [18], a commonly used method for PMR-based kinship inference. Specifically, PMR cutoffs for degree classification are determined after normalizing PMRs by the median PMR; the normalization step attempts to account for variation in background relatedness. Additionally, third-degree kinship classification is limited to sample pairs whose “expected mismatches” exceed 3,000. Finally, `fastpmr` supports two output modes: CSV files or a compressed `numpy` archive (`.npz`).

## Supporting information

Supplementary Table S1: AADR v66.p1 Potential Spurious Duplicates

Supplementary Table S2: AADR v66.p1 Potential Cryptic Duplicates (Annotated)

Supplementary Table S3: AADR v66.p1 Potential Cross-Site Relatives

Supplementary Table S4: Chichen Itza Recomputed PMRs

Supplementary Table S5: AADR v62.0.p1 Potential Spurious Duplicates

Supplementary Table S6: AADR v62.0.p1 Potential Cryptic Duplicates

Supplementary Table S7: AADR v62.0.p1 Potential Cross-Site Relatives

## Declaration

### Ethics approval and consent to participate

No ethics approval is required. This work re-analyzes public, previously published aDNA data. Several of our findings reported in Results: Quality control and discovery at scale in the global aDNA record may revise published interpretations of ancient individuals. We report these findings to improve the accuracy of the shared aDNA record and have aimed to describe them with care. Before submission of this work, we contacted the corresponding authors of the affected studies and invited them to comment on our re-analysis.

### Consent for publication

Not applicable.

## Availability of data and materials

Code and documentation for `fastpmr` are freely available at https://github.com/ehuangc/fastpmr. The version of `fastpmr` (v0.2.4) and associated data used in this work are archived on Zenodo [39]. No new genomic data suitable for downstream analysis were generated in this work. Sequencing of the Twist-captured BSB004 library, performed as a diagnostic check, showed that most reads derived from a KLK002 library (Results: Quality control and discovery at scale in the global aDNA record); we therefore do not deposit these reads. Other genomic data used in this work are available from their respective publications [1, 23, 25, 28, 29, 31–34].

## Authors’ contributions

E.C.H. developed the software, ran the computational experiments, created the visualizations, and wrote and edited the manuscript. H.R. contributed research directions, supervised the work, and reviewed and edited the manuscript.

## Acknowledgements

We gratefully acknowledge David Reich for helpful discussion regarding the AADR and Lyazzat Musralina for assistance in investigating the data from BSB004 and KLK002. We also thank Florence Pautet for sharing her implementation of the `hapROH` analysis with a custom haplotype reference panel.

