## Supplementary Information for "Systematic inference of biological relatedness across the global ancient DNA record"

### Supplementary Information for
### Systematic inference of biological relatedness across the global ancient DNA record

Edward C. Huang[1,*] and Harald Ringbauer[1]

[1]Department of Archaeogenetics, Max Planck Institute for Evolutionary Anthropology, Leipzig, Germany

#### Contents

Supplementary Note 1: Gaussian process analysis pg. S1

Supplementary Note 2: ROH analysis of samples from medieval England pg. S3

Supplementary Note 3: Radiocarbon dating of samples from Kazakhstan pg. S5

Additional Supplementary Figures pg. S6

#### Supplementary Note 1: Gaussian process analysis

To expand our AADR PMR analysis beyond within-locality sample pairs, we fit a custom Gaussian process model to a random subset of European (and adjacent) pairs in the AADR, including between-locality sample pairs. This Gaussian process model predicts PMRs for hypothetical sample pairs given input spatial coordinates and times. It represents a PMR as a sum of three terms: two per-sample latent effects tied to each sample's location and time, and a within-pair effect depending only on the spatial and temporal separation between the two samples. Then, predicting the PMR for a spatially and temporally coincident sample pair probes the per-sample PMR component at a single point, which we interpret as local genetic diversity.

We spatially interpolate AADR PMR data at various time points by using the Gaussian process model to predict these "coincident PMRs" (Figure S1). Notably, we observe a hotspot in southeastern Europe at 1000 BP, associated with higher coincident PMRs and potentially higher local genetic diversity. This hotspot could reflect admixture during early-medieval migrations in the region [1]. We note that this Gaussian process analysis aims to highlight the scientific potential of large-scale ancient relatedness data, and more work is necessary to develop robust inference methods for such data. For example, Gaussian processes are poorly suited to representing strict discontinuities, so this analysis may obscure migration barriers and other discontinuous population genetic patterns. Additionally, we find that the final plots are somewhat sensitive to the choice of training parameters, so we further caution against overinterpretation of this analysis.

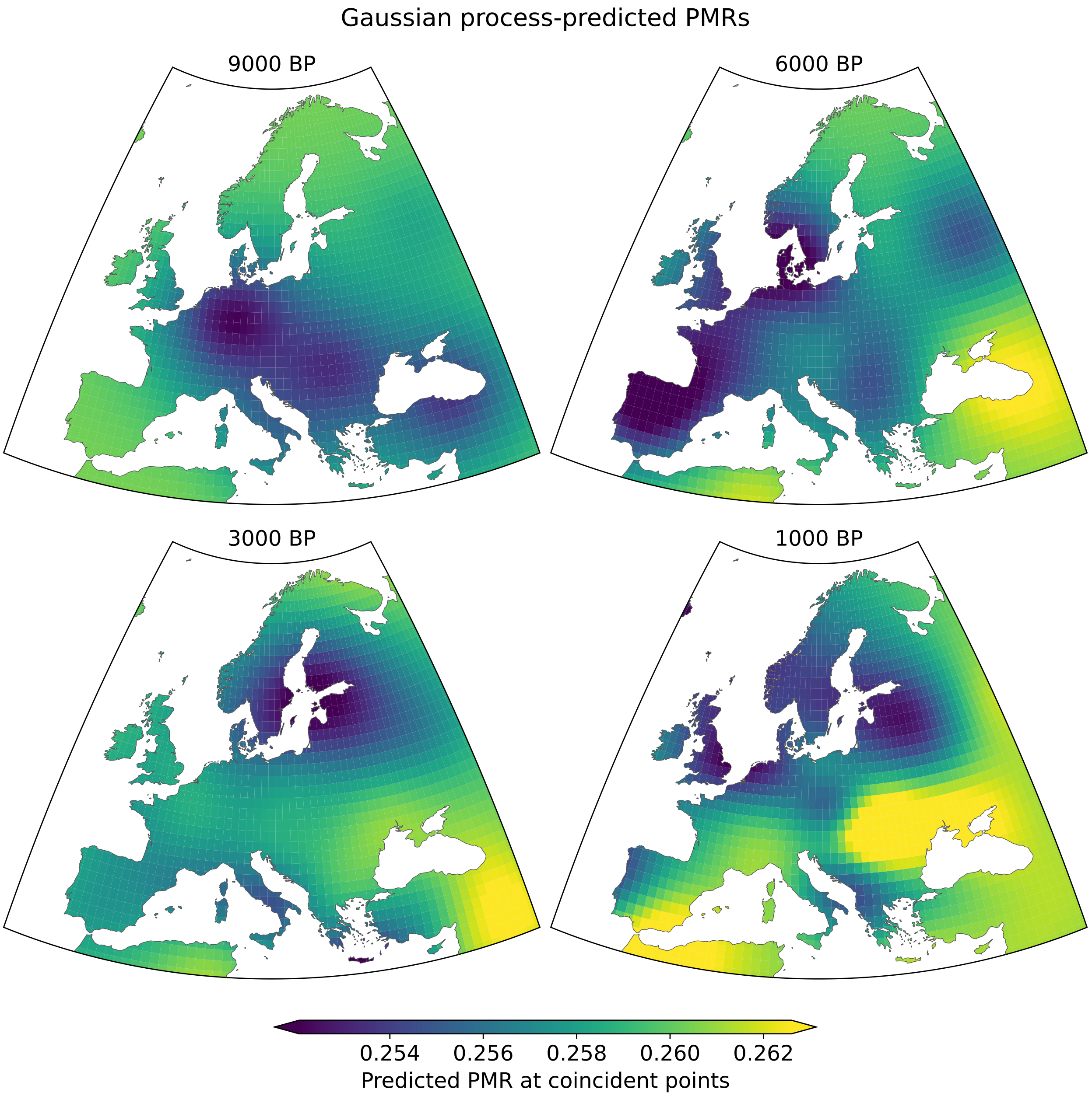


Figure S1: Gaussian process-predicted PMRs of spatially and temporally coincident sample pairs, plotted at 1° latitude/longitude grid points and various time points. To avoid bias from close relatives, sample pairs with a PMR more than five standard deviations below the mean are not included in the training data.

### Supplementary Note 2: ROH analysis of samples from medieval England

We estimate runs of homozygosity (ROH) for the three potential genetic duplicates from medieval England — all inferred to have Ashkenazi Jewish-like ancestry (see Results: Quality control and discovery at scale in the global aDNA record). To do so, we run the software hapROH [2], which is designed for calling ROH segments in human ancient DNA, with a custom reference panel of 1000 Genomes haplotypes restricted to all biallelic SNPs with minor allele frequency $> 5\%$[3]. Initial testing indicates that this configuration enables accurate detection of ROH segments >4 centimorgans long for samples with as low as $\sim$0.1x average whole-genome sequencing coverage [in preparation]. As such, all three target samples are well within the application range.

The inferred ROH patterns for the three target samples are very similar. For example, each sample shares almost-identical long ROH segments on chromosomes 3, 7, 13, and 19. These results would be extremely unlikely if the samples were not genetically identical. Under Mendelian inheritance, even one pair of full siblings only co-inherits an ROH segment with probability $\leq 0.25$, as each parent transmits one of the two possible haplotypes at random.

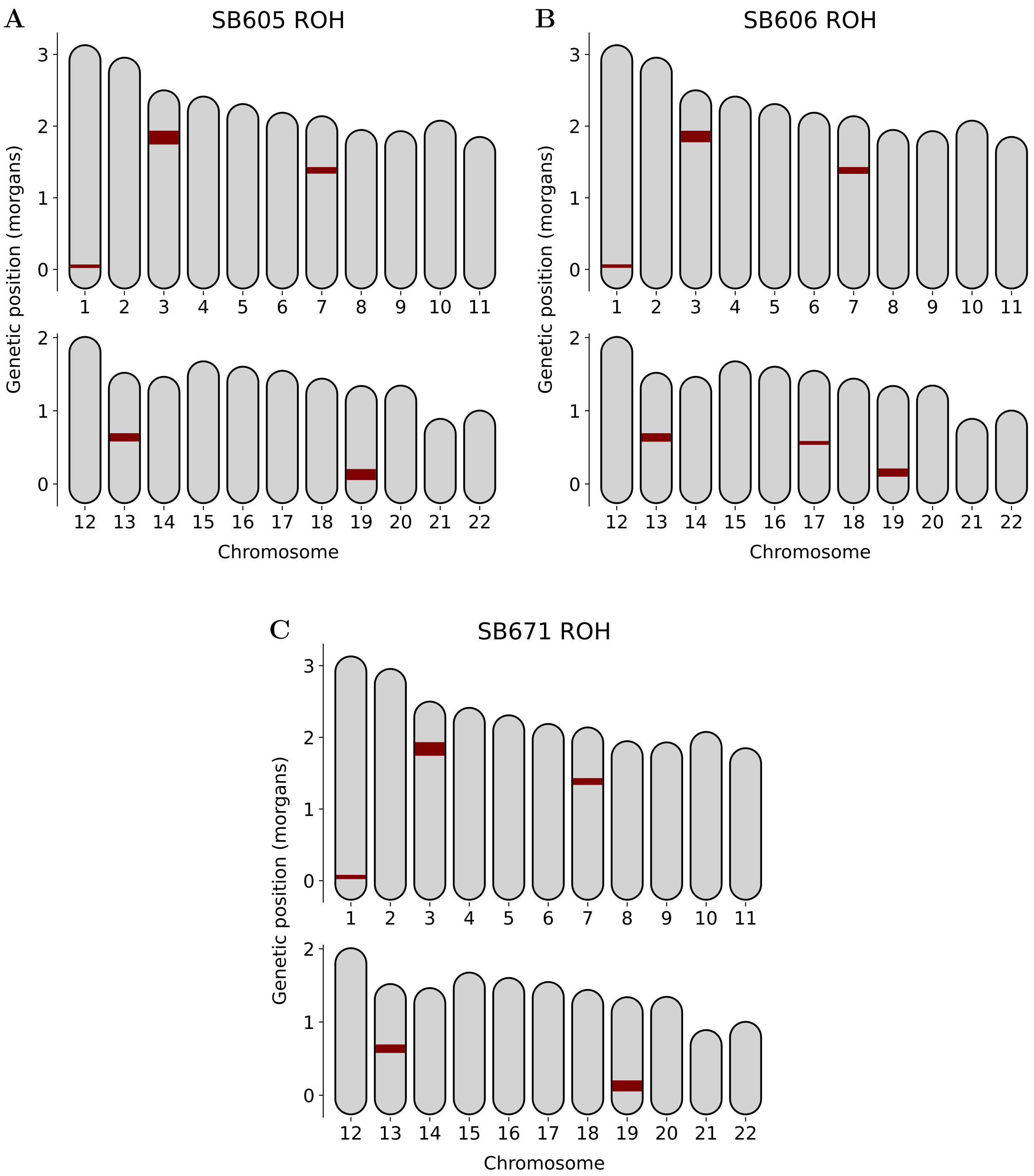


Figure S2: Inferred ROH for the three potential cryptic duplicates from Medieval England. ROH segments > 4 centimorgans long are depicted on the 22 autosomal chromosomes.

### Supplementary Note 3: Radiocarbon dating of samples from Kazakhstan

One tooth from each of BSB004 and KLK002 was dated by AMS at the Curt-Engelhorn-Zentrum Archäometrie (CEZA), Mannheim and calibrated with IntCal20 [4] in OxCal 4.4 [5]. BSB004 (MAMS-87639) yielded $2477 \pm 18$ BP, calibrated to 764–537 cal BC (93.5%) and 532–517 cal BC (2.0%) at 95.4% probability. KLK002 (MAMS-87640) yielded $923 \pm 17$ BP, calibrated to 1040–1169 cal AD (95.4%). Both samples had C:N ratios within the range for well-preserved collagen.

#### Additional Supplementary Figures

A

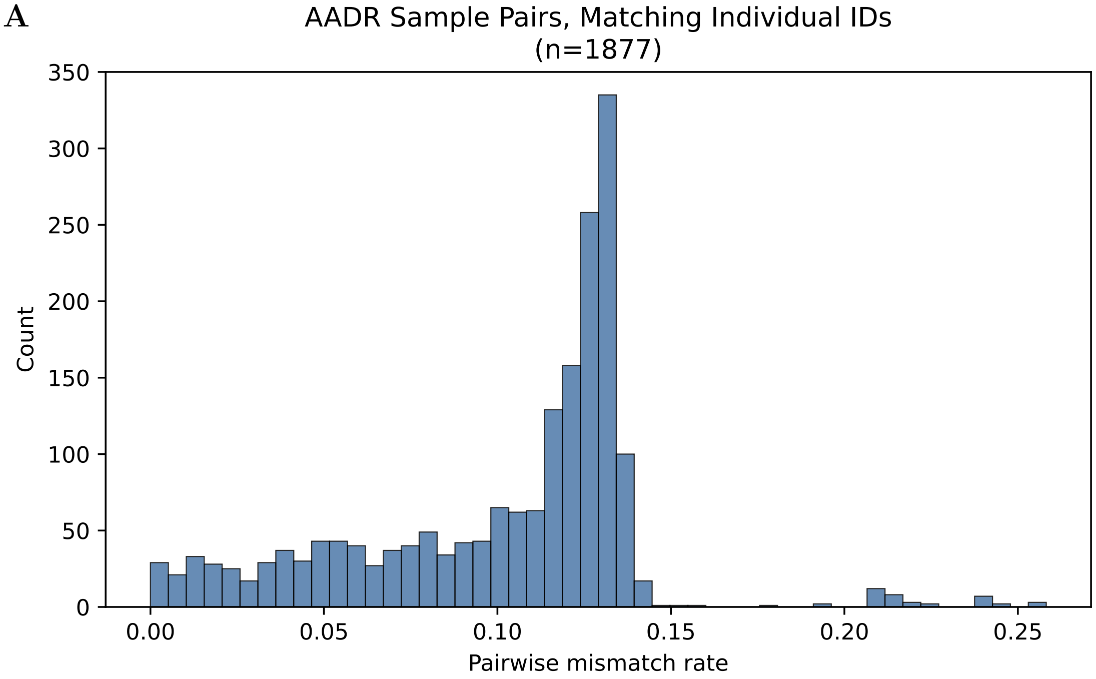


B

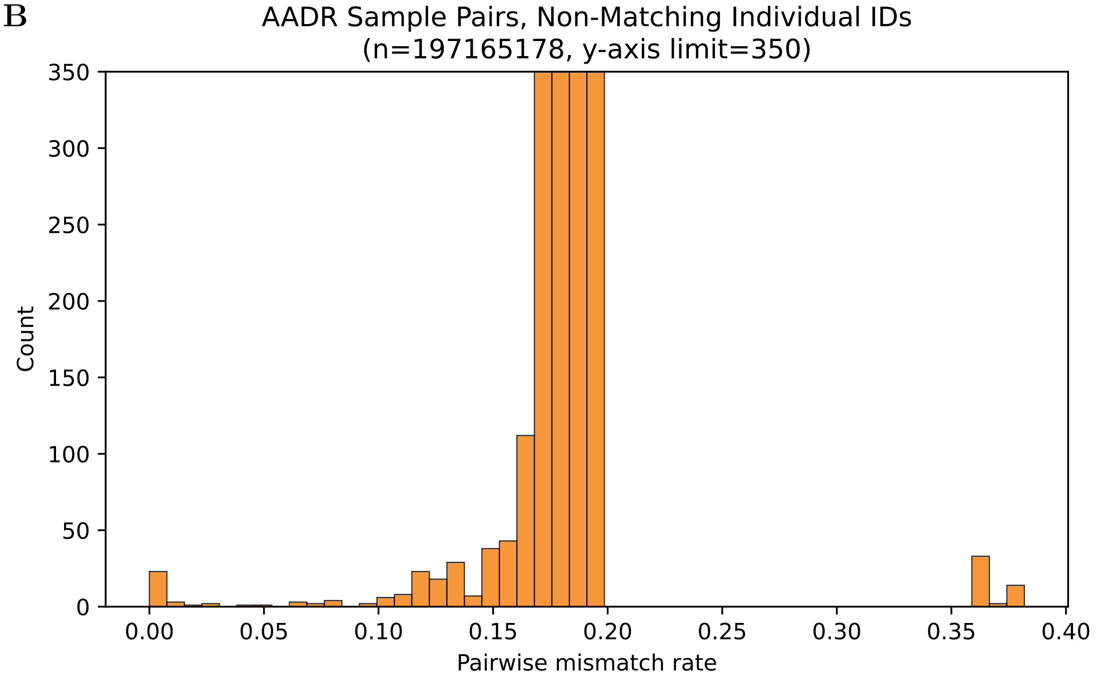


Figure S3: PMR distributions of AADR sample pairs with (**A**) matching and (**B**) non-matching individual IDs. The y-axis limit in (**B**) is set to 350.
